# Efficient inactivation of African swine fever virus by a highly complexed iodine combined with compound organic acids

**DOI:** 10.1101/2022.01.10.475769

**Authors:** Mengnan Qi, Li Pan, Ying Gao, Miao Li, Yanjin Wang, Lian-Feng Li, Chen Ji, Yuan Sun, Hua-Ji Qiu

## Abstract

African swine fever (ASF) is a highly contagious disease with high morbidity and mortality caused by African swine fever virus (ASFV). Cleaning and disinfection remain one of the most effective biosecurity measures to prevent and control the spread of ASFV. In this study, we evaluated the inactivation effects of highly complexed iodine (HPCI) combined with compound organic acids (COAs) against ASFV under different conditions. The results showed that the inactivation rates of the disinfectants on the reporter ASFV increased in dose- and time-dependent manners, the best inactivation effects were obtained when the compatibility ratio of HPCI and COAs was 5:1 at 25°C. Furthermore, there were no significant differences by comparing the efficacy of HPCI combined with COAs (HPCI+COAs) in inactivating wild-type ASFV and the reporter ASFV (*P* > 0.05). ASFV of 10^4.0^ TCID_50_/mL was completely inactivated by 0.13% HPCI (0.0065% effective iodine), 0.06% COAs or 0.13% HPCI+COAs (approximately 0.0054% effective iodine), respectively, while 10^6.0^ TCID_50_/mL ASFV was completely inactivated by 1.00% HPCI (0.05% effective iodine), 0.50% COAs or 1.00% HPCI+COAs (0.042% effective iodine), respectively. Therefore, HPCI+COAs had synergistic effects to inactivate ASFV. This study demonstrated that HPCI+COAs could rapidly and efficiently inactivate ASFV and represent an effective compound disinfectant for the control of ASF.

**IMPORTANCE:** African swine fever (ASF) is a highly contagious disease with high morbidity and mortality caused by African swine fever virus (ASFV). Due to the lack of commercial vaccines and treatment available for ASF, effective disinfectants and the proper use of them are extremely essential to inactivate ASFV. The significance of this research is in searching for an ideal disinfectant that not only has the advantages of low toxicity and non-pollution but also can inactivate ASFV rapidly and efficiently. In this study, we proved that HPCI+COAs not only exhibited low cytotoxicity, but also could completely inactivate ASFV within 5 min at 4°C, 25°C and 37°C. In addition, HPCI+COAs had synergistic effects on inactivated ASFV. Thus, HPCI +COAs could be used as an effective disinfectant for the control of ASF.

## INTRODUCTION

African swine fever (ASF) caused by African swine fever virus (ASFV) is a highly contagious hemorrhagic viral disease with high morbidity and mortality (1, 2). The World Organization for Animal Health (OIE) lists ASF as a notifiable animal disease. ASFV infects both domestic pigs and wild boar, and is responsible for serious economic and production losses (2, 3). ASFV is a member of the *Asfivirus* genus within the *Asfarviridae* family, which has a linear double-stranded DNA genome of 170–194 kb encoding 150–167 proteins, including structural and host immunomodulatory proteins (4, 5). The ASFV virion consists of an internal core, an internal lipid membrane, an icosahedral capsid, and an outer lipid envelope (6, 7). ASFV is relatively stable and can survive in meat and blood for several months at room temperature. The disease can be transmitted through various routes, including vehicles, personnel exchange and direct or indirect contact (8).

Since the first description in 1921, ASF has been circulating in different parts of the world, which is endemic in Africa and has caused recurrent outbreaks in other territories (4). Since August 2018, a highly virulent genotype II ASFV has been spread to China and successively to Mongolia, Vietnam, Cambodia, Laos, North Korea, Philippines, Myanmar, South Korea, Indonesia, and other Asia-Pacific countries (9, 10). In China, small-scale pig farms with weak biosecurity systems account for a large proportion, which may be one reason for the rapid spread of ASF in China (8).

Due to the lack of commercial vaccines and treatment available for ASF, the prevention and control of ASF is mainly based on quarantine, culling and strict biosecurity measures (11). Effective disinfectants and the proper use of them are extremely essential to inactivate ASFV. The ideal disinfectant should be characterized by: high efficiency, low toxicity and non-pollution (12, 13). Chemicals such as glutaraldehyde, sodium hydroxide, phenols, and quaternary ammonium salts can be used to prevent and control ASF, yet these disinfectants are irritating or corrosive (14, 15). In addition, sodium hypochlorite can effectively inactivate ASFV, whereas organic matter negates the effects (16, 17). In the field, the viruses on pig farms are encased in organic matter, which can make disinfectants ineffective. At present, the main using methods of disinfections include drinking water disinfection, environmental disinfection and animal in vitro disinfection, which can reduce the probability of swine infected with ASFV while irritating the porcine skin, mucous membrane, and cannot prevent swine detoxing through the way such as oral cavity or effectively inactivating ASFV that exist in the upper gastrointestinal tract (18). Therefore, it is necessary to develop a safe disinfectant that can be used not only for environmental disinfection but also for disinfection *in vitro* and *in vivo*.

Highly complexed iodine (HPCI) is a high polyiodine-containing disinfectant with a high safety profile. Different from traditional iodine preparations, HPCI has the advantages of high stability, and oral safety. The mechanism of iodine-inactivating pathogens is that free iodine can combine with the hydrocarbon, sulfhydryl, amino and hydroxyl groups on the protein amino acid chain, leading to protein denaturation and precipitation (19, 20). Compound organic acids (COAs) are the acidifiers composed of fatty acids such as formic acid, propionic acid and lactic acid. COAs can release H^+^ and hydrolyze nucleic acid or proteins by permeating into the biofilm of pathogens (21). In addition, COAs have the ability to enhance body immunity due to their inhibiting the basic metabolism of bacteria and regulating pH in bacteria (22).

As commercial feed additives, COAs possess good palatability while the price of HPCI is higher than other commonly used disinfectants. Thus, the use of the combination of HPCI and COAs is expected to decrease costs and can be added to drinking water or feed. Moreover, both HPCI and COAs have the advantages of non-toxicity and non-pollution, which can inactivate ASFV effectively. In view of the advantages above and the urgent need for the prevention and control of ASF, we evaluated the inactivation effects of HPCI combined with COAs (HPCI+COAs) on ASFV.

## RESULTS

### Cytotoxicity assessment of HPCI+COAs in HEK293T and PAM cells

HEK293T cells and PAMs were inoculated with HPCI+COAs at different concentrations and cultured for 8 h. The cell viability was detected by CCK-8 kit. The results showed that when the mixing ratio of HPCI and COAs was 1:10, 1:5, 1:2, 1:1, 2:1, 5:1 and 10:1, the disinfectant was determined to have less cytotoxicity against HEK293T cells than PAMs. The cell viability on HEK293T cells or PAMs was above 80% when the concentration of the mixture was less than 1.00% or 0.25%, respectively (Fig. S1).

### Neutralization assessment

The purpose of the neutralizer identification test was to determine whether the neutralizer could neutralize the residual HPCI or COAs. Firstly, the cytotoxicity of neutralizing products was evaluated in HEK293T cells and PAMs. The results showed that the neutralizing products had no effect on HEK293T cells and PAM cells (Fig. S2). The viral titers in treatment groups 3 to 5 were close to 4.5 log10, indicating that the neutralizer had no significant effect on the replication of ASFV and could neutralize HPCI and COAs (Tables 1 and 2).

**Table. 1.** Effects of the neutralizer on HPCI.

| Group | Treatment | ASFV titer ($\log_{10}$ TCID<sub>50</sub>/mL) |
| --- | --- | --- |
| 1 | (HPCI + ASFV) + Medium | $-\infty$ |
| 2 | (HPCI + ASFV) + Neutralizer | $-\infty$ |
| 3 | (Neutralizer + ASFV) + Medium | $4.25 \pm 0.35$ |
| 4 | (HPCI + Neutralizer) + ASFV | $4.50 \pm 0.00$ |
| 5 | ASFV + Medium | $4.46 \pm 0.06$ |
| 6 | Medium only | $-\infty$ |

$-\infty$ : ASFV was undetectable.

**Table 2.** Effects of the neutralizer on COAs.

| Group | Treatment | ASFV titer ($\log_{10}$ TCID<sub>50</sub>/mL) |
| --- | --- | --- |
| 1 | (COAs + ASFV) + Medium | $-\infty$ |
| 2 | (COAs + ASFV) + Neutralizer | $-\infty$ |
| 3 | (Neutralizer + ASFV) + Medium | $4.50 \pm 0.10$ |
| 4 | (COAs + Neutralizer) + ASFV | $4.46 \pm 0.05$ |
| 5 | ASFV + Medium | $4.54 \pm 0.05$ |
| 6 | Medium only | $-\infty$ |

$-\infty$ : ASFV was undetectable.

### The optimal ratio of HPCI and COAs

To determine the best compatibility ratio of HPCI and COAs, the inactivated effects of the cocktails mixing in different proportions were evaluated. Different concentrations of HPCI+COAs were interacted with 10^4.0^-10^6.0^ TCID_50_/mL reporter ASFV, and incubated for 30 min at room temperature (25°C), and then stopped by the neutralizer. The virus inactivation rates were calculated by infectivity assays on HEK293T cells. The inactivation rates of the cocktails on the reporter ASFV increased in a dose-dependent manner, and the better inactivation effect was achieved when the compatibility ratio of HPCI and COAs was 5:1 (Fig. 1).

**Fig. 1.**
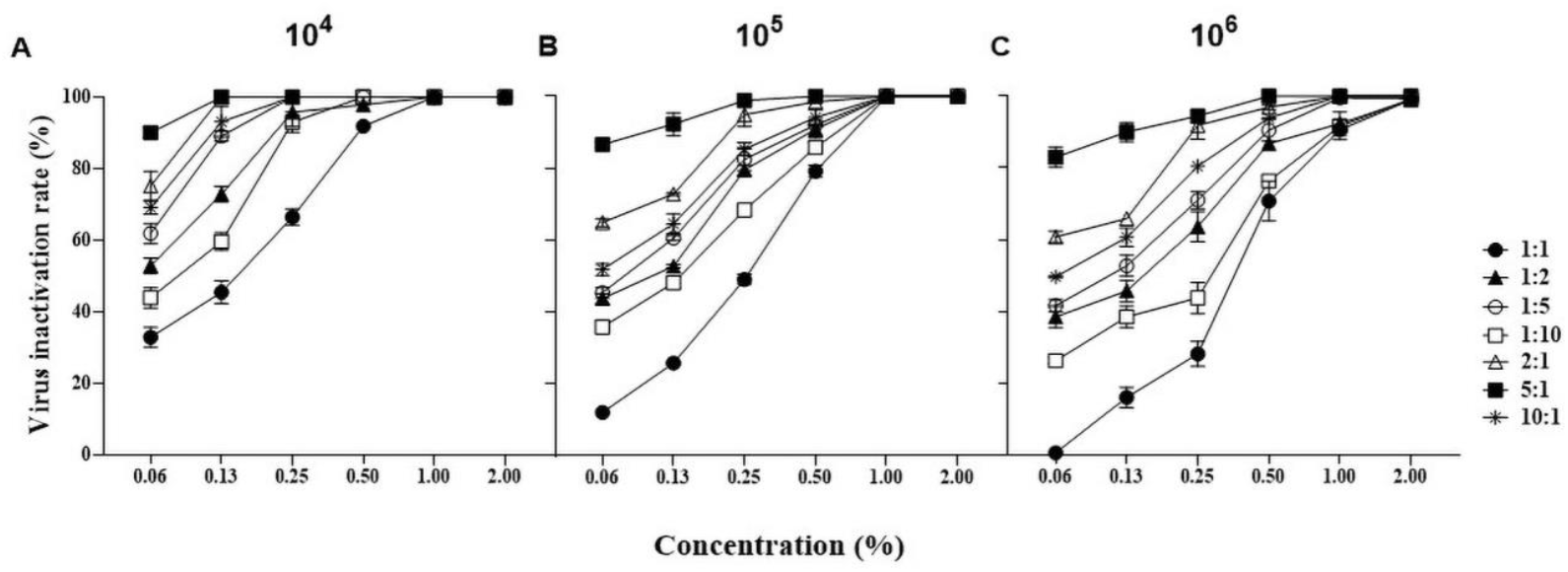
Inactivation of the reporter ASFV by HPCI+COAs. 10^4.0^, 10^5.0^ and 10^6.0^ TCID_50_/mL reporter ASFV were inactivated with HPCI+COAs at 25°C for 30 min, respectively, and stopped by the neutralizer and the resulting virus inactivation rates were calculated by infectivity assay on HEK293T cells.

### The inactivation effects were time-dependent

Different concentrations of disinfectants were interacted with 10^6.0^ TCID_50_/mL reporter ASFV and incubated for 5, 15 and 30 min, respectively, and then stopped by the neutralizer and the virus inactivation rates were determined by infectivity assays on HEK293T cells. 100% inactivation rates of the reporter ASFV were observed when inactivated with HPCI, COAs or HPCI+COAs as lower as 1.00% within 30 min, indicating that these three disinfectants can inactivate ASFV rapidly and efficiently (Fig. 2). In addition, the inactivation rates were the lowest at 5 min while the highest at 30 min after treating the reporter ASFV with different concentrations of three disinfectants, indicating that the inactivated effects were time-dependent (Fig. S3).

**Fig. 2.**
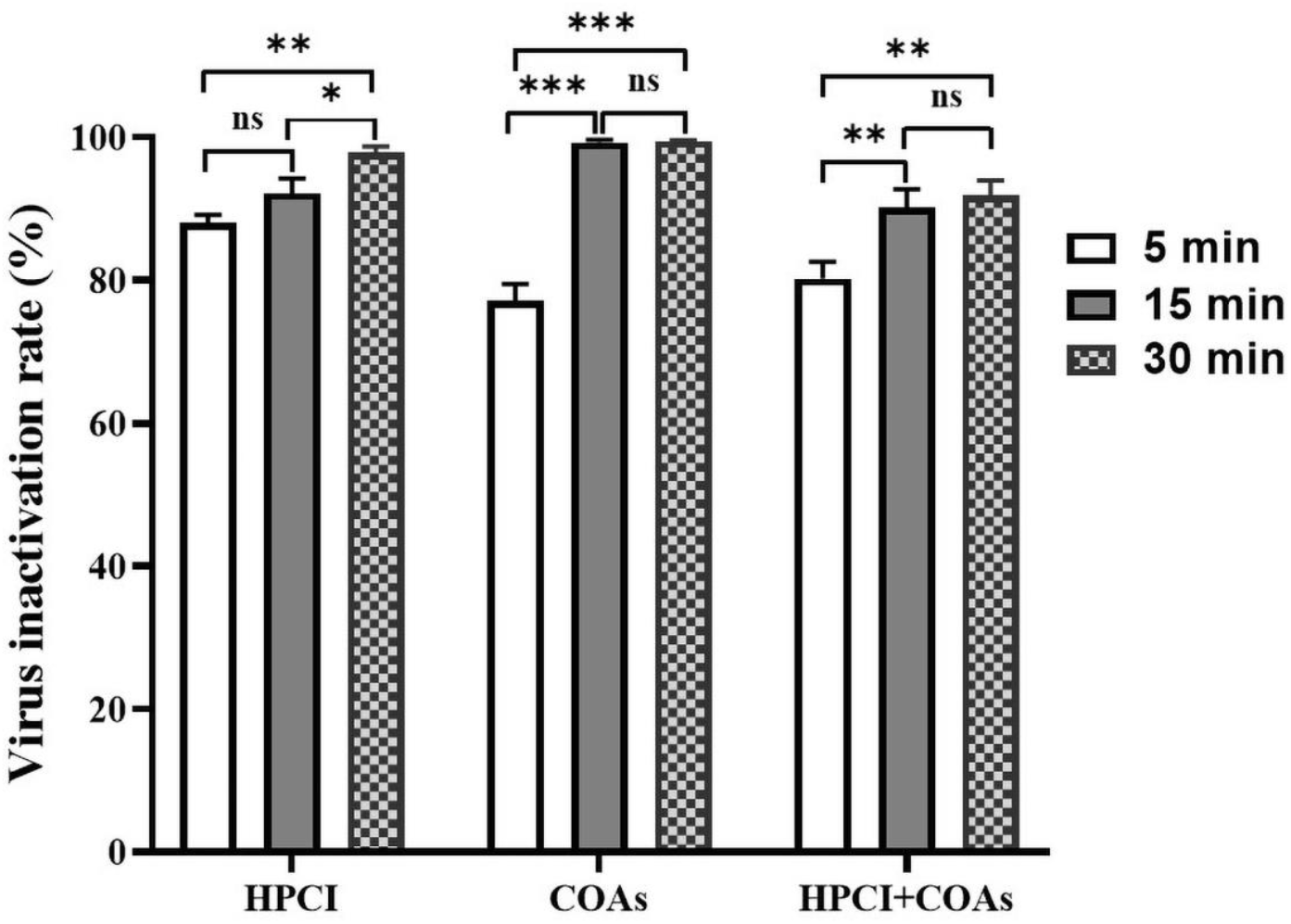
Inactivation of the reporter ASFV by HPCI+COAs in different times. 10^6.0^ TCID_50_/mL reporter ASFV was inactivated with 1.00% HPCI, COAs or HPCI+COAs for 5, 15 or 30 min, and stopped by the neutralizer and the resulting virus inactivation rates were calculated by infectivity assay on HEK293T cells.

### The optimal temperature for inactivating ASFV

Different concentrations of disinfectants were applied to 10^6.0^ TCID_50_/mL reporter ASFV and incubated at different temperatures for 5 min, and then stopped by the neutralizer and the virus inactivation rates were determined by infectivity assays on HEK293T cells. It was important to mention that no significant differences were found in the inactivation rates of the reporter ASFV between 4°C and 37°C when inactivated with HPCI, HPCI+COAs as lower as 1.00% (Fig. 3). However, the inactivation rate of the reporter ASFV at 25°C was better than those of 4°C and 37°C, indicating that 25°C was the optimal temperature for HPCI and COAs to inactivate the reporter ASFV (Fig. S4).

**Fig. 3.**
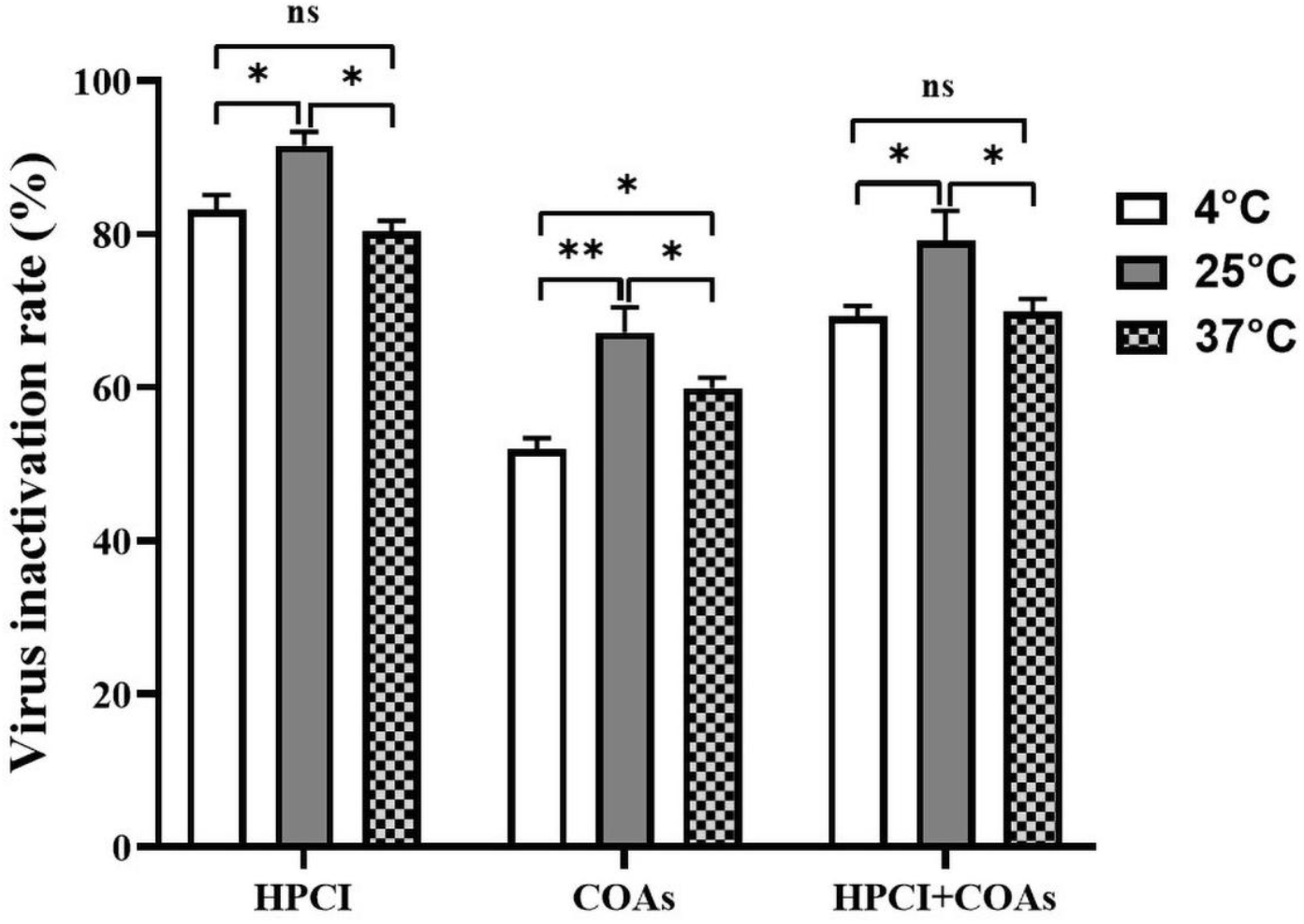
Inactivation of the reporter ASFV by HPCI+COAs at different temperatures. 10^6.0^ TCID_50_/mL reporter ASFV was inactivated with 1.00% HPCI, COAs or HPCI+COAs at 4, 25 or 37°C for 5 min, and stopped by the neutralizer and the resulting virus inactivation rates were calculated by infectivity assay on HEK293T cells.

### Bovine serum albumin attenuates inactivation effects of disinfectants

The disinfectants diluted with PBS containing different concentrations of bovine serum albumin (BSA) were applied to 10^6.0^ TCID_50_/mL reporter ASFV, respectively, and incubated at 25°C for 30 min. Then the mixtures were inoculated into cell plates to evaluate the effect of the disinfectants under the interference of organic matter. The results showed that 0.3% BSA had no significant effect on the inactivation effects of the reporter ASFV, whereas, 3% BSA had a significant effect, indicating that the high concentration of organic matter had an effect on the inactivation effects of disinfectants to ASFV (Fig. 4).

**Fig. 4.**
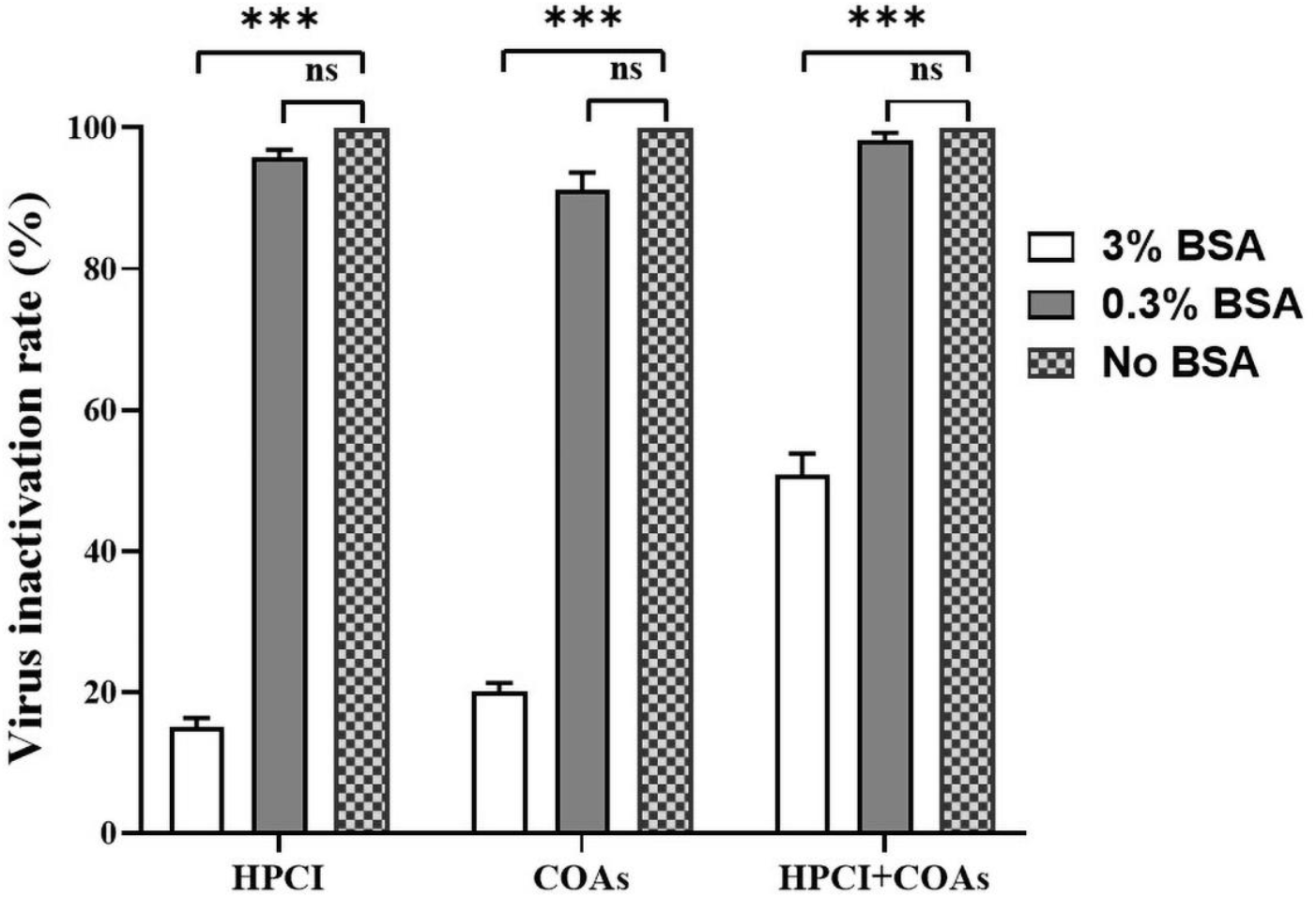
Inactivation of the reporter ASFV interfered with different concentrations of BSA by HPCI+COAs. 10^6.0^ TCID_50_/mL reporter ASFV was inactivated with HPCI, COAs or HPCI+COAs containing no BSA, 0.3% or 3% BSA for 30 min, and stopped by the neutralizer and the resulting virus inactivation rates were calculated by infectivity assay on HEK293T cells.

### ASFV could be inactivated by disinfectants effectively

Different concentrations of disinfectants were applied to 10^4.0^-10^6.0^ TCID_50_/mL wild-type ASFV Pig/HLJ/18 strain (WT-ASFV) (GenBank: MK333180.1) or reporter ASFV and incubated for 30 min at 25°C, respectively, and then stopped by the neutralizer and the virus inactivation rates were determined by infectivity assays on PAMs or HEK293T cells. The results showed that 100% inactivation rates of either WT-ASFV or the reporter ASFV were observed when inactivated with HPCI or HPCI+COAs as lower as 1.00%, and inactivated with COAs as lower as 0.50% (Table 3). Moreover, 10^4.0^ TCID_50_/mL WT-ASFV or reporter ASFV could be inactivated completely with 0.13% HPCI, 0.06% COAs or 0.13% HPCI+COAs while 0.50% HPCI, 0.13% COAs or 0.50% HPCI+COAs could completely inactivate 10^5.0^ TCID_50_/mL ASFV (Tables S1 and S2). There were no significant differences by HPCI+COAs in inactivating WT-ASFV and the reporter ASFV (*P* > 0.05). Taken together, all disinfectants can effectively and rapidly inactivate ASFV.

**Table 3.** Inactivation (%) of 10^6.0^ TCID_50_/mL ASFV by disinfectants.

| Disinfectant | Virus | Concentration of disinfectants (%) |  |  |  |  |  |
| --- | --- | --- | --- | --- | --- | --- | --- |
|  |  | 0.06 | 0.13 | 0.25 | 0.50 | 1.00 | 2.00 |
| HPCI | Reporter ASFV | 90.07 ± 0.05 | 95.90 ± 0.03 | 98.94 ± 0.01 | 99.98 ± 0.00 | 100.00 ± 0.00 | 100.00 ± 0.00 |
|  | WT-ASFV | 97.85 ± 0.03 | 98.17 ± 0.02 | 99.94 ± 0.00 | 99.95 ± 0.00 | 100.00 ± 0.00 | 100.00 ± 0.00 |
| COAs | Reporter ASFV | 99.97 ± 0.00 | 99.98 ± 0.00 | 99.99 ± 0.00 | 100.00 ± 0.00 | 100.00 ± 0.00 | 100.00 ± 0.00 |
|  | WT-ASFV | 99.94 ± 0.00 | 99.97 ± 0.00 | 99.99 ± 0.00 | 100.00 ± 0.00 | 100.00 ± 0.00 | 100.00 ± 0.00 |
| HPCI+COAs | Reporter ASFV | 83.00 ± 0.03 | 90.00 ± 0.03 | 94.49 ± 0.00 | 99.99 ± 0.00 | 100.00 ± 0.00 | 100.00 ± 0.00 |
|  | WT-ASFV | 90.00 ± 0.01 | 93.19 ± 0.04 | 95.36 ± 0.06 | 99.85 ± 0.00 | 100.00 ± 0.00 | 100.00 ± 0.00 |

### Verification of ASFV inactivation in HEK293T cells or PAMs

To confirm whether ASFV has lost its infectivity after being inactivated by 1.00% HPCI, COAs or HPCI+COAs, the disinfected ASFVs were collected and passaged in HEK293T cells or PAMs. Green fluorescence of the reporter ASFV was observed by inverted fluorescence microscope while immunofluorescence assay (IFA) and hemadsorption test (HAD) were used to determine the inactivation effects of WT-ASFV. The results indicated that no live ASFVs were detected (Fig. 5 and Fig. S5).

**Fig. 5.**
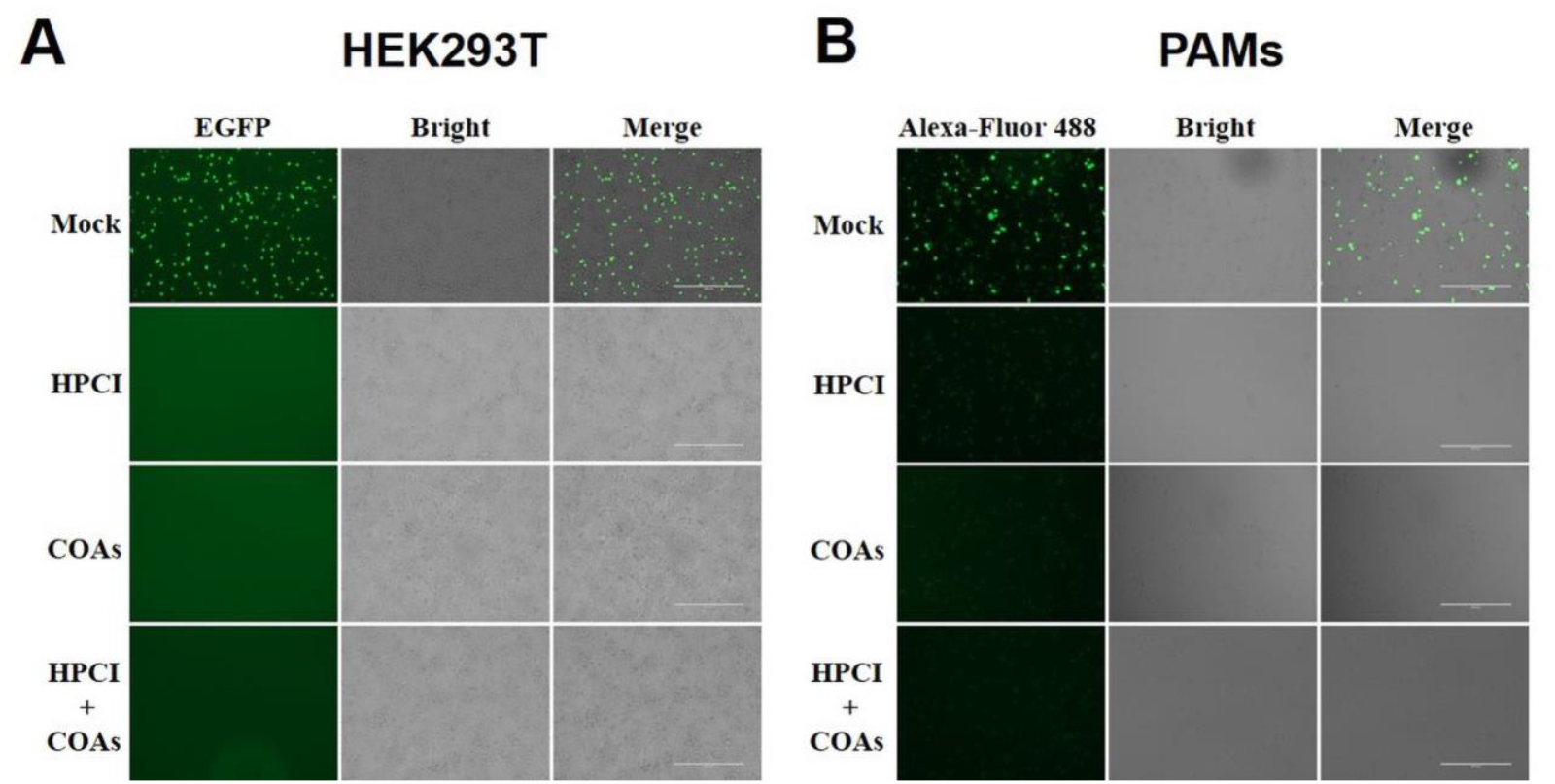
Verification of inactivation of ASFV by HPCI+COAs. ASFV was treated with HPCI, COAs or HPCI+COAs, and then collected and inoculated into HEK293T cells (A) or PAMs (B) for repeated infection tests. Green fluorescence of the reporter ASFV was observed by inverted fluorescence microscope while immunofluorescence assay (IFA) was used to determine the inactivation effects of WT-ASFV.

## DISCUSSION

The continuous spread of ASF has caused huge economic losses to the pig industries in China (23). As no available commercial vaccines or treatable drugs, strict biosecurity measures have become one of the effective measures to prevent and control ASF. Effective disinfectants are necessary for this issue. HPCI and COAs can efficiently inactivate pathogenic microorganisms by oxidizing properties and will be applicable in various scenarios. Therefore, the efficacy of HPCI+COAs in inactivating ASFV was evaluated under different conditions in this study.

HPCI, compared with other iodine disinfectants, is more stable and has no stimulation and damages to cells and mucosal tissues, which can ensure that HPCI will not damage the oral cavity, nasal cavity, digestive tract and respiratory mucosa of animals. Previous studies have shown that ASFV could be inactivated by immersion or spray with 0.25–5% HPCI in 5-30 min (24). ASFV has a relatively high pH resistance range that the actual survival time of the virus is relatively short when the pH value is lower than 3.4 (2). Therefore, acid disinfectants can be selected to inactivate ASFV. High-quality organic acids can not only inactivate ASFV but also have nutritional value and immunomodulatory effects (25).

In this study, we evaluated the inactivation of ASFV by HPCI+COAs. Our data showed that the better inactivation effect was achieved when the mixture ratio of HPCI and COAs was 5:1. In addition, the better temperature for HPCI and COAs to inactivate the reporter ASFV was 25°C and the inactivation effects were time-dependent. Moreover, the results showed that HPCI+COAs had the same efficacy against WT-ASFV and the reporter ASFV, which had synergistic effects on inactivation of ASFV.

ASFV can be transmitted by direct or indirect contact between infected animals, infected pig products, or infected fomites, such as transport vehicles (26, 27). Organic materials such as soil and manure can interfere with disinfectants (HPCI and COAs) (28). Based on the above results, we recommend that the ground and equipment of pig farms were thoroughly washed and then disinfected. It is suggested that 0.13% HPCI, 0.06% COAs and 0.13% HPCI+COAs can be used in the non-epidemic areas. And in the epidemic area, 1.00% HPCI, 0.50% COAs and 1.00% HPCI+COAs should be used to inactivate ASFV.

In conclusion, HPCI+COAs can be used as an effective disinfectant for the control of ASF.

## MATERIALS AND METHODS

### Preparation of cells and viruses

Primary porcine alveolar macrophages (PAMs) were prepared from 3-week-old specific-pathogen-free (SPF) pigs and maintained in RPMI 1640 medium (Gibco, USA) supplemented with 10% fetal bovine serum (FBS) (Gibco, USA), 200 mg/mL streptomycin and 200 IU/mL penicillin. Human embryonic kidney 293T cells (HEK293T cells) were maintained in RPMI 1640 medium (Gibco, USA) supplemented with 10% fetal bovine serum (FBS) (Gibco, USA). PAMs or HEK293T cells (10^6.0^ cells/mL) were seeded into 6-well plates with 2 mL per well or 96-well plates with 100 μL per well and incubated at 37°C in a humidified incubator with 5% CO_2_. The follow-up tests were carried out after cell adhesion.

WT-ASFV was inoculated into PAMs (6-well plates) and incubated at 37°C for 72 to 96 h. The reporter ASFV that can infect HEK293T cells and express the green fluorescent protein (ASFV-P60-CD2v-EGFP) was constructed by our lab in Harbin Veterinary Research Institute (HVRI) of the Chinese Academy of Agricultural Science (CAAS), and was inoculated into HEK293T cells (6-well plates) and incubated at 37°C for 72 to 96 h. After three freeze-thaw cycles, the cell cultures were centrifuged at 12,000 ×*g* for 5 min to remove cells and other debris and the supernatants with virus suspension was stored at −80°C until use. The viral titer was determined by the Reed and Muench method (29). All experiments with live ASFV in this study were performed in the biosafety level 3 (BSL-3) facilities in HVRI of the CAAS.

### Preparation of commercial disinfectants

HPCI containing 5% effective iodine (China Patent Application No. 202110654174.3) was contained in the form of a tan liquid from Guangdong Ruiji Biotechnology Co., Ltd., China, and COAs (China Patent Application No. 20190823035705) were contained in the form of an orange liquid from Beijing Huijiayuan Animal Husbandry Tech Co., Ltd., China. According to the manufacturer’s descriptions, highly complexed iodine is the main component of HPCI, and other components mainly promote fusion and improve the stability of HPCI. COAs are acidifiers composed of short- and medium-chain fatty acids, which have rapid disinfection, strong stability and surface adhesion ability. Both HPCI and COAs can be stably stored for at least two years at 21-27°C.

### Cytotoxicity test

According to the instructions, HPCI and COAs were aseptically filtered with 0.22-μm filters and mixed in different proportions and diluted to different concentrations with sterile hard water. Each dilution of the disinfectants was added to the 96-well cell culture plate with HEK293T cells or PAMs prearranged, and each dilution was performed in triplicates. The control group was added with the same amount of sterile hard water. All the cell plates were cultured in an incubator with 5% CO_2_ at 37°C for 8 h. The morphology of cells treated with different disinfectants was observed by the inverted microscope, and then the activity of these treatments on the cells was determined using Cell Counting Kit-8 (CCK-8) (APExBIO, K1018) according to the manufacturer’s instructions and the technical specification for disinfection.

### Identification test of neutralizer

To evaluate whether the neutralizer could neutralize the residual disinfectant, the reporter ASFV was incubated at room temperature with HPCI or COAs for 30 min, followed by the addition of neutralizer (RPMI 1640 medium containing 5 g/L sodium thiosulfate and 10% FBS) (Group 1 in Tables 1 and 2) or RPMI 1640 medium (Group 2) (1:9, v/v) (26). After incubation for 30 min at room temperature, HEK293T cells (in 96-well plates) were inoculated with 100 μL mixtures and incubated for 4 days at 37°C incubator with 5% CO_2_. The same procedures were used for Groups 3 to 5 as shown in Tables 1 and 2. The inactivation effects of HPCI or COAs on the reporter ASFV were evaluated by infectivity assays on HEK293T cells and the viral titers were determined as described previously. Average values and standard deviations of three independent experiments were calculated.

### Virucidal assay

The reporter ASFV or WT-ASFV (10^4.0^, 10^5.0^ and 10^6.0^ TCID_50_/mL) were incubated with different concentrations of the disinfectants (0.06-2.00%), then HEK293T cells or PAMs (in 96-well plates) were inoculated with the mixtures of virus and disinfectant, respectively, and incubated at 37°C with 5% CO_2_. Three replicates were performed at each concentration, plus positive control, different concentrations of solution control and blank cell control. After 8-h incubation, the mixture was replaced with the fresh cell culture medium and cell viability was observed at 3 to 5 days post-inoculation (dpi). The supernatant fluid with virus suspension was stored at −80°C until use. The viral titer of the supernatant fluid was determined.

To confirm whether ASFV has lost its infectivity after being inactivated, the viral supernatants were harvested and re-inoculated onto fresh HEK293T cells or PAMs to culture for five days. The mixtures were passaged three times in HEK293T cells or PAMs. IFA and HAD were used to determine the inactivation effects. These methods were performed as described previously (30).

### Statistical analysis

Statistical analyses were performed using GraphPad Prism 8 software (GraphPad Software Inc., USA). Differences between groups were examined for statistical significance using Student’s *t*-test. An unadjusted *P*-value of less than 0.05 was considered significant.

## ACKNOWLEDGMENTS

This study was supported by the Natural Science Foundation of China (U20A2060 and 32072854) and the Natural Science Foundation of Heilongjiang Province of China (JQ2020C002).

